# BiGCN: Leveraging Cell and Gene Similarities for Single-cell Transcriptome Imputation with Bi-Graph Convolutional Networks

**DOI:** 10.1101/2024.04.05.588342

**Authors:** Yoshitaka Inoue, Ethan Kulman, Rui Kuang

## Abstract

**Motivation:** RNA profiling at the single-cell level is essential for characterizing the molecular activities and functions of individual cells. The current technical limitations of single-cell RNA sequencing (scRNA-seq) technologies can lead to a phenomenon known as “dropout”, where a significant portion of gene expression is not captured. Dropout is particularly prominent in genes with low or sparse expression, greatly impacting the reliability and interpretability of scRNA-seq data. Consequently, various techniques have been developed to estimate missing gene expression using imputation, often by either modeling similarities in gene expression among cells or using gene co-expression, but rarely both.

**Results:** In this study, we introduce a Bi-Graph Convolutional Network (BiGCN), a deep learning method that leverages both cell similarities and gene co-expression to capture cell-type-specific gene co-expression patterns for imputing scRNA-seq data. BiGCN constructs both a cell similarity graph and a gene co-expression graph, and employs them for convolutional smoothing in a dual two-layer Graph Convolutional Networks (GCNs). The embeddings from the two GCNs can subsequently be combined to facilitate the final imputation. BiGCN demonstrates superior performance compared to state-of-the-art imputation methods on both real and simulated scRNA-seq data. Additionally, BiGCN outperforms existing methods when tasked with clustering cells into cell types. We also perform a novel validation using a PBMC scRNA-seq dataset, and this experiment supports that BiGCN’s imputations are more realistic than competing imputation methods. In both the imputation and the cluster tasks, BiGCN consistently outperformed two variants of BiGCN that solely relied on either the gene co-expression graph or cell similarity graph. This indicates that the two graphs offer complimentary information for imputation and cell clustering, underscoring the importance of incorporating both types of information.

**Code Availability:** https://github.com/inoue0426/scBiGCN.

## Introduction

Single-cell RNA sequencing (scRNA-seq) technologies have revolutionized gene expression studies at the single-cell level [Tang et al., 2009] by enabling researchers to analyze the whole transcriptome in individual cells. This approach offers a more accurate and detailed view of gene expression patterns in terms of their diversity and complexity within a population of cells. By analyzing gene expression patterns at the single-cell level, researchers can gain insights into the roles and interactions of different cell types, which contributes to a better understanding of complex tissues and organs.

However, analyzing scRNA-seq data poses challenges due to dropout events [Kharchenko et al., 2014]. Dropout can occur due to a method’s limited sensitivity, or the challenge in detecting lowly expressed genes. To address this issue, various imputation methods have been developed Hou et al. [2020]. While many of these methods rely on the similarity of gene expression across different cells to learn a *cell-cell similarity graph* [Tran et al., 2021, Dijk et al., 2018], only a few incorporate gene co-expression [Rao et al., 2021]. In many cases, cells can be clustered using only on marker genes [Ianevski et al., 2022], although genes can exhibit significant variation in expression levels among cells [Haque et al., 2017]. Therefore, both cell similarities and gene co-expression are crucial structures to model for more accurate and robust scRNA-seq data imputation.

This study introduces a bi-directional method called BiGCN (Bi-Graph Convolutional Networks), which leverages cell similarity and gene co-expression to recover scRNA-seq dropouts. BiGCN combines the embeddings learned from two Graph Convolutional Networks [Kipf and Welling, 2016], one representing a cell similarity graph and the other a gene co-expression graph. These embeddings are then used for imputing scRNA-seq data. An overview of the BiGNN architecture is given in Fig. 1 (explained in detail in section 3).

**Fig. 1.**
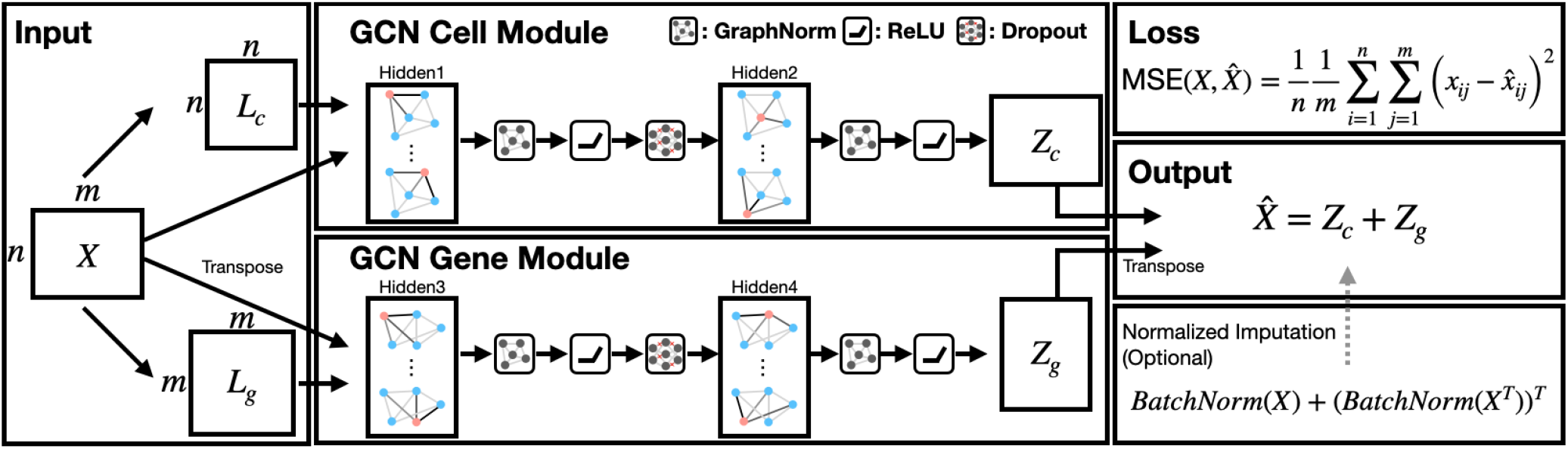
Overview of BiGCN. First, a GCN, Graph Norm, ReLU, and Dropout layers are applied to the adjacency matrix of cells *L*_*c*_ and genes *L*_*g*_ with the original matrix *X*. The output is then reshaped to match the size of the original matrix using GCN, Graph Norm, and ReLU, resulting in *Z*_*c*_ ∈ ℝ^*n×m*^ and *Z*_*g*_ ∈ ℝ^*n×m*^. These matrices are combined to obtain 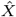. Both 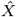 and *X* are inputs for the MSELoss function. Additionally, a normalized imputation option enhances stability by adding batch-normalized data to the imputation results.

## Related work

Existing scRNA-seq data imputation methods can be categorized into three main groups: deep learning-based, affinity-based, and statistics-based.

In deep learning-based approaches, a subset of scRNA is often used as the training data. Deep Neural Networks (DNNs), Graph Neural Networks (GNNs), and other techniques are employed with a loss function such as zero inflated negative binomial loss (ZINBLoss) or mean square error (MSELoss) to train on this subset. Afterward, imputation is performed on the remaining data. Examples of these methods include scGNN [Wang et al., 2021], DeepImpute [Arisdakessian et al., 2019], GNNImpute [Xu et al., 2021a], and DCA [Eraslan et al., 2019]. Affinity-based imputation methods use the similarity between cells to infer missing values. These methods typically identify cells with similar gene expression profiles and use the gene expression data from those cells to fill in missing values in other cells. Examples of affinity-based imputation methods include MAGIC [Dijk et al., 2018], scImpute [Li and Li, 2018], kNN-smoothing [Wagner et al., 2018], and DrImpute [Gong et al., 2018].

Statistical imputation methods use statistical models to estimate missing values. These methods can account for gene correlations and large amounts of missing data. Examples of statistical-based imputation methods include ALRA [Linderman et al., 2022], AdImpute [Xu et al., 2021b], SDImpute [Qi et al., 2021], and scISR [Tran et al., 2022].

DeepImpute [Arisdakessian et al., 2019], GNNImpute [Xu et al., 2021a], MAGIC [Dijk et al., 2018], and ALRA [Linderman et al., 2022] are widely used among these methods. These are used as baselines in this paper. In addition, we also utilized BiRW [Xie et al., 2015] as a baseline for scRNA imputation to show the importance of both networks and dealing with a cold-start problem.

## Method

In this section, we formulate the scRNA imputation problem as graph embedding learning problem, introduce the network architecture of BiGCN, and introduce a variation of BiGCN suitable for normalized imputation with parameterized cell-wise and gene-wise rescalings.

### Constructing affinity matrix

Given a log transformed (read count plus one) gene expression matrix *X* ∈ ℝ^*n×m*^, where *n* represents the number of rows (cells) and *m* represents the number of columns (genes), the linear kernel defined below is used to construct the affinity matrix.

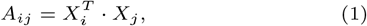

where *A* is an affinity matrix, and *X*_*i*_ and *X*_*j*_ represent the rows containing the gene expression data for cells *i* and *j*, respectively. Since *X* consists entirely of non-negative values and Equation 1 defines a linear function of the rows of *X*, the entries of *A* will also be non-negative. To improve the stability of the model we opted for a sparse graph structure. We utilized only the top 15% of the affinity matrix *A*, denoted as *Â*, defined as follows:

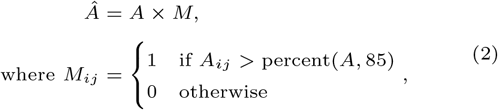

where percent(*A, p*) returns a threshold value at rank 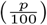 among the entries in *A*. Then, the following normalization of the Laplacian matrix *Â* is performed to obtain *L*,

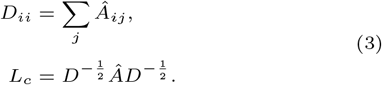

We applied the above procedure cell-wise over *X* ∈ ℝ^*n×m*^ to obtain the matrix *L*_*c*_ and similarly gene-wise over *X*^*T*^ ∈ ℝ^*m×n*^, to get the matrix *L*_*g*_. These matrices were then used as inputs for BiGCN as affinity matrices.

### Bi-Graph Convolutional Network

Bi-Graph Convolutional Network (BiGCN) includes two separate graph convolution network modules for convolution over the cell and gene graphs, respectively. Fig 1 illustrates the overall architecture of BiGCN. Cell graph *L*_*c*_, gene graph *L*_*g*_ and expression data matrix *X* are the inputs for the two GCN modules. Specifically, (*L*_*c*_,*X*) and (*L*_*g*_,*X*^*T*^) are embedded by each GCN module and then decoded to 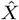 and 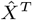, which are summed together to construct the imputed matrix 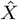. The mean squared error between 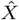 and *X* are used as the loss function. It is also possible to generate a normalized imputation by incorporating normalized original data to stabilize the training process. This variant will be explained in the following section. In the following, we define the functions for the cell module and the definitions for the gene module can be similarly defined. The formula of the first layer in the cell module is as follows:

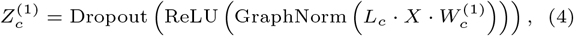

where 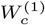 is the weight matrix for the first layer. Here we utilized GraphNorm [Cai et al., 2021] defined as follows:

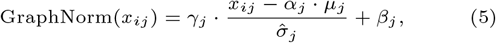

where *x* is an input for GraphNorm, 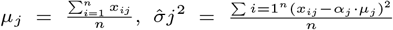, and *γ*_*j*_, *β*_*j*_ are the affine parameters as commonly applied in other normalization methods.

Then, 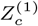 is the input of the fourth layer through another series of operations.

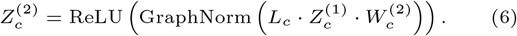

The same structure is applied to the gene GCN module with (*L*_*g*_, *X*^*T*^) as inputs in Equation 4 to train the weight matrix *W* ^(1)^ and obtain the output 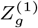. Then, 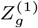 is the input to the structure in Equation 6 with *L*_*g*_ and weight matrix 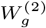 to obtain the output 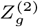. Finally, the imputed matrix 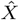 in obtained by adding the outputs of the two GCN modules together, The final result of the imputation is obtained by adding these matrices together,

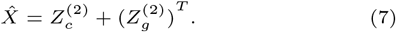

MSE loss function was minimized between the original data matrix *X* and the imputed data matrix 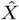.

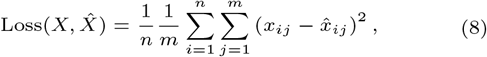

where *x*_*ij*_ and 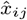 are the entry at row *i* and column *j* in *X* and 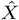, respectively.

In our study, we optimized the MSELoss using Adam [Kingma and Ba, 2014], an adaptive optimization algorithm. Adam dynamically adjusts the learning rate based on the estimated second moment of the gradients for faster convergence.

### Normalized imputation with parameterized rescaling

To perform better downstream analysis, such as clustering, a variant of GCN is also introduced to obtain a normalized imputation 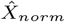. This imputation captures the cell-specific and gene-specific scales by adjustment with parameterized batch-normalization:

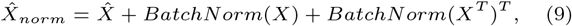

where 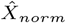 is utilized in the MSE loss function. Here, batch normalization (BatchNorm) [Ioffe and Szegedy, 2015] is defined as follows:

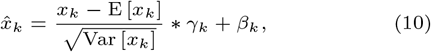

where 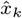 is row *k* in BatchNorm(X) and *γ*_*k*_ and *β*_*k*_ are learnable parameters.

### Parameter tuning

We utilized Optuna [Akiba et al., 2019], a Python library for automatic hyperparameter optimization, to conduct hyperparameter tuning for the imputation and clustering experiments. It optimized 16 parameters, including six dropout rates, six hidden layers, and additional parameters such as epochs, learning rate, and optimizer. We explored various parameter settings to evaluate the performance of the model. We tested values ranging from 0.1 to 0.5 for the dropout. The hidden layer sizes varied from 2 to 1024. The number of epochs was set between 10 and 500. We searched for the learning rate within the range of 1e-5 to 1e-1. Additionally, we experimented with three different optimizers: Adam, RMSprop, and SGD.

To optimize the MSE values, we optimized the hyper-parameters with the 10%-dropout data and used the same parameters for all other simulation datasets of larger dropout ratios. For the clustering tasks, we used the same set of parameters for the ten annotated scRNA-seq datasets. These parameters are shown in Table 1.

**Table 1.** Optimal parameters.

| Params | $\log(\text{read\_counts} + 1)$ | Normalized expressions | Type | Tuning Range | Description |
| --- | --- | --- | --- | --- | --- |
| dropout1 | 0.3 | 0.2 | Float | [0.1, 0.5] | Dropout rate for layer 1 |
| dropout2 | 0.1 | 0.2 | Float | [0.1, 0.5] | Dropout rate for layer 2 |
| hidden0 | # of Cell | # of Cell | Int | None | Number of cells for the initial layer |
| hidden1 | # of Gene | # of Gene | Int | None | Number of genes for the initial layer |
| hidden2 | 128 | 256 | Int | [128, 1024] | Number of hidden units for layer 2 |
| hidden3 | 1024 | 512 | Int | [128, 1024] | Number of hidden units for layer 3 |
| epochs | 1500 | 100 | Int | [50, 3000] | Number of training epochs |
| learning rate | 0.0001 | 0.01 | Float | [1e-5, 1e-1] | Learning rate for the optimizer |
| optimizer | Adam | Adam | Categorical | Adam, SGD, RMSprop | Optimization algorithm |

### GCNs with gene module or cell module

To gain further insight into the role of gene co-expression and cell similarities, the two GCN modules: GCN Cell and GCN Gene, can be also applied separately for imputation as shown in Figure 1. GCN Cell performs graph convolution using only the cell similarity network. GCN Gene performs graph convolution using the gene co-expression network. Comparing these two variants with BiGCN can reveal the impact that cell similarity, gene co-expression, and their combination have on imputation and clustering. This becomes particularly relevant when the number of cells and genes varies in scRNA-seq data.

### Data preparation

#### Simulation data for imputation

We used scDesign2 [Sun et al., 2021] to create simulated scRNA-seq data. The base input data was the mouse intestinal epithelial tissue [Haber et al., 2017], GSE accession number GSE92332, from scDesign2. After fitting scDesign2 with the base inputs, we used it to generate simulated data with no dropout. We first generated a dataset of 1000 samples and 15,962 genes divided into 8 clusters, denoted as “1k cell simulation data” in the results. Artificial dropouts were then randomly introduced into the simulation data. The dropout rate was incrementally set at intervals of 10%, ranging from 10%-90%. A 0% dropout (original data) was used as the ground truth. Additionally, we sampled a second dataset comprising 5,000 cells and 5,000 genes, and a third dataset comprising 10,000 samples and 5,000 genes. Note that the 5,000 genes selected for each dataset were those with the largest variances.

These additional datasets are denoted as “5k cell simulation data” and “10k cell simulation data”.

#### PBMC scRNAseq data with aggregated bulk reference

We also created an *aggregated bulk reference* data set derived from real scRNA-seq data to further evaluate imputation performance in a realistic setting. We acquired a large 68k PBMC dataset from the 10x Genomics data repository. This dataset contains 68,579 cells with 32,738 genes profiled by the Visium platform. We randomly partitioned the data into two subsets: 10,000 cells to undergo imputation, and the remaining 58,579 cells to derive an aggregate bulk reference. We chose to keep only genes that are expressed in at least 10 cells of the 10,000 that will undergo imputation, leaving 12,016 genes.

Next, we computed the correlations between the 10,000 cells set aside for imputation, and the other 58,579 cells. The mean correlation was 0.775 with a standard deviation of 0.107. For each cell in the 10,000 cells set aside for imputation, we found its *k*-nearest neighbors in the other 58,579 cells (for *k* = 10, 20, 30), and the average gene expression across these *k*-nearest neighbors is used as an aggregated bulk reference. For each *k*, this results in an *aggregate bulk reference matrix* with 10,000 rows, and 12,016 columns. These averaged matrices were labeled as the ‘10-neighborhood average,’ ‘20-neighborhood average,’ and ‘30-neighborhood average,’ respectively. Note that if the number of nearest neighbors are less than the top-*k* needed for aggregation, all the nearest neighbors are kept.

#### Single-cell RNA-seq data for clustering

Ten scRNAseq datasets were used to evaluate a method’s ability to cluster cells into cell types, and these datasets are listed in Table 2. The ten datasets were obtained from various human tissues: Human Endometrial assembloids (HEA), Human Adipose Tissue (HAT), Human Duodenal Organoids (HDO), Human Cell Atlas bone marrow (HBM), Human Brain (HB), Human Embryo (HE), Human Mesenchymal Progenitor Cell (HMPC), Human Ventral Midbrain (HEVM), Human Induced Pluripotent Stem Cells (HIPSC), and Human Liver Transplantation (HLT). These datasets cover a range of number of cell types (4 to 26), cell counts (1014 to 12000), gene counts (10960 to 28643), and gene densities (2.36% to 30.27%). HEA, HB, HE, HEVM, HIPSC, and HLT were collected from Gene Expression Omnibus [Edgar et al., 2002]. The accession numbers are GSE168405, GSE118257, GSE156456, GSE76381, GSE129096, and GSE189539 respectively. HAT, HDO, and HMPC were downloaded from Single Cell PORTAL (https://singlecell.broadinstitute.org/single_cell). The accession numbers are SCP1903, SCP1318 and SCP1027. HBM was obtained from Seurat [Hao et al., 2021].

**Table 2.** scRNAseq Datasets.

| Dataset | Tissue Type | n_class | n_cell | n_gene | density [%] | Reference |
| --- | --- | --- | --- | --- | --- | --- |
| HIPSC | Human Induced Pluripotent Stem Cells | 5 | 7445 | 28643 | 2.36 | [Selewa et al., 2020] |
| HB | Human Brain | 8 | 8000 | 21581 | 5.02 | [Jäkel et al., 2019] |
| HBM | Human Cell Atlas bone marrow | 8 | 12000 | 17369 | 5.80 | [Hay et al., 2018] |
| HLT | Human Liver Transplantation | 14 | 8196 | 19988 | 6.94 | [Wang et al., 2022b] |
| HAT | Human Adipose Tissue | 10 | 9082 | 27202 | 6.95 | [Strieder-Barboza et al., 2022] |
| HEA | Human Endometrial assembloids | 11 | 5829 | 20697 | 8.04 | [Rawlings et al., 2021] |
| HEVM | Human Ventral Midbrain | 26 | 1977 | 19531 | 11.76 | [La Manno et al., 2016] |
| HE | Human Embryo | 8 | 1014 | 25742 | 24.38 | [Wang et al., 2022a] |
| HDO | Human Duodenal Organoids | 9 | 2484 | 14679 | 26.74 | [Mead et al., 2022] |
| HMPC | Human Mesenchymal Progenitor Cell | 4 | 6656 | 10960 | 30.27 | [Loureiro et al., 2022] |

The n\_class, n\_cell, and n\_gene describe the number of classes, cells, and genes, respectively.

#### Data preprocessing

A preprocessing step is conducted where genes with no expression are excluded, followed by a logarithmic transformation of counts plus 1. However, in processing for DeepImpute, which uses raw data as input, imputation is carried out without the logarithmic transformation, and the transformation is applied to the output.

When assessing clustering, we exclusively considered genes expressed in more than 5% of the cells, and similarly, we only included cells with expression data for more than 5% of genes.. Following this step, logarithmic, library size, and square root transformations were applied.

### Evaluation and visualization

We utilize Mean Squared Error (MSE) and Correlation to evaluate our imputation results. When assessing clustering, we also considered the Adjusted Rand Index (ARI). In the context of our research, it is desirable to have a low MSE, and a high Correlation and ARI. We kept the parameters for the baseline methods the same as those optimized in their original papers.

UMAP (Uniform Manifold Approximation and Projection) [McInnes et al., 2018] was used to visualize high-dimensional data by projecting it into a lower-dimensional space using graph topological embedding. Another commonly used dimensionality reduction technique for visualization is t-SNE (t-Distributed Stochastic Neighbor Embedding) Van der Maaten and Hinton [2008]. In this study, we utilized the Scanpy package [Wolf et al., 2018] to perform UMAP and t-SNE. We have included two representative UMAP visualizations in the main text and the remaining ones in the Appendix.

## Results

### Imputation results of simulation data

In this section, we use simulation data with a known ground truth to evaluate BiGCN’s imputation performance against GCN Cell, GCN Gene, ALRA, DeepImpute, MAGIC, and BiRW. This simulation data has from 10% to 90% of its entries randomly impacted by dropout events.

The results in Table 3 show the MSE for the simulation data with 1000 cells and 15962 genes. Here we show the results for 10%, 40%, 50%, 60%, 70%, and 80% dropouts. BiGCN always has the lowest MSE, followed by GCN Gene and GCN Cell. This indicates that integration of the similarity between cells and genes does provide additional benefits for imputation. Furthermore, the results in Table 4 show the correlation coefficients. BiGCN achieves the highest correlation in four datasets and the second highest for two. While MAGIC showed the highest correlation in one experiment with the very low drop-out rate 10%, which is very similar to the ground-truth data before introducing dropouts. Thus, it has a low indication of the actual imputation performance.

**Table 3.** MSE on 1k cell simulation data (red, orange and blue indicate the top three methods)

| Dropout Ratio | 10% | 40% | 50% | 60% | 70% | 80% |
| --- | --- | --- | --- | --- | --- | --- |
| Density | 19.93 | 13.29 | 11.08 | 8.86 | 6.64 | 4.43 |
| ALRA | 0.9152 | 1.5823 | 1.7641 | 2.3058 | 2.0073 | 3.3598 |
| MAGIC | 0.1650<sup>◇</sup> | 0.2145 | 0.2407 | 0.2713 | 0.3060 | 0.3420 |
| DeepImpute | 0.5934 | 0.5226 | 0.4596 | 0.4463 | 0.3863 | 0.3613 |
| GNNImpute | 0.3390 | 0.3438 | 0.3454 | 0.3503 | 0.3568 | 0.3675 |
| BiRW | 0.2341 | 0.3314 | 0.3543 | 0.3705 | 0.3817 | 0.3914 |
| BiGCN | 0.1644<sup>★</sup> | 0.2005<sup>★</sup> | 0.2204<sup>★</sup> | 0.2474<sup>★</sup> | 0.2783<sup>★</sup> | 0.3162<sup>★</sup> |
| GCN Cell | 0.1738 | 0.2137<sup>★</sup> | 0.2376<sup>★</sup> | 0.2618<sup>★</sup> | 0.2903<sup>★</sup> | 0.3277<sup>★</sup> |
| GCN Gene | 0.1672<sup>★</sup> | 0.2018<sup>◇</sup> | 0.2241<sup>◇</sup> | 0.2475<sup>◇</sup> | 0.2826<sup>◇</sup> | 0.3164<sup>◇</sup> |

**Table 4.** Correlations on 1k cell simulation data (red, orange and blue indicate the top three methods).

| Dropout Ratio | 10% | 40% | 50% | 60% | 70% | 80% |
| --- | --- | --- | --- | --- | --- | --- |
| Density | 19.93 | 13.29 | 11.08 | 8.86 | 6.64 | 4.43 |
| ALRA | 0.4447 | 0.4151 | 0.4144 | 0.4009 | 0.4087 | 0.3958 |
| MAGIC | 0.7128<sup>★</sup> | 0.6933<sup>★</sup> | 0.6834<sup>★</sup> | 0.6706<sup>★</sup> | 0.6511<sup>★</sup> | 0.6197<sup>★</sup> |
| DeepImpute | 0.4371 | 0.4425 | 0.4560 | 0.4629 | 0.4786 | 0.4839 |
| GNNImpute | 0.2488 | 0.2270 | 0.2265 | 0.2163 | 0.2067 | 0.1811 |
| BiRW | 0.6267 | 0.5787 | 0.5481 | 0.4977 | 0.4114 | 0.2967 |
| BiGCN | 0.7075<sup>◇</sup> | 0.6991<sup>★</sup> | 0.6952<sup>★</sup> | 0.6896<sup>★</sup> | 0.6787<sup>★</sup> | 0.6219<sup>◇</sup> |
| GCN Cell | 0.6881 | 0.6644 | 0.6552 | 0.6443 | 0.6295 | 0.6123 |
| GCN Gene | 0.7025 | 0.6980<sup>◇</sup> | 0.6940<sup>◇</sup> | 0.6887<sup>◇</sup> | 0.6783<sup>◇</sup> | 0.6528<sup>★</sup> |

Next, we conducted the same imputation evaluation on the 5k cell simulation data. Table 5 shows the MSE for the 5k cell simulation data. In four datasets, BiGCN consistently achieved the lowest MSEs, and for two datasets, it ranked second only to GCN Cell and GCN Gene. Notably, when working with 1k cell data, GCN Gene performed better, while for datasets with the same cell-gene ratio, GCN Cell performed better. Furthermore, Table 6 displays the correlation results. BiGCN exhibited the best performance for three datasets, ranked second for two datasets and third for one dataset, MAGIC attained the second-best results, followed by GCN Cell in the third position. Again, MAGIC generated slightly better correlation in the datasets with lower dropout rates but performs poorly on the sparser datasets.

**Table 5.** MSE on 5k cell simulation data (red, orange and blue indicate the top three methods).

| Dropout Ratio | 10% | 40% | 50% | 60% | 70% | 80% |
| --- | --- | --- | --- | --- | --- | --- |
| Density | 30.37 | 20.25 | 16.88 | 13.51 | 10.13 | 6.75 |
| ALRA | 2.1954 | 3.1238 | 3.7172 | 4.4207 | 5.5927 | 7.2328 |
| MAGIC | 0.2667<sup>◇</sup> | 0.3671 | 0.4220 | 0.4857 | 0.5549 | 0.6252 |
| DeepImpute | 0.9721 | 0.8337 | 0.8383 | 0.7649 | 0.6323 | 0.5764<sup>★</sup> |
| GNNImpute | 0.6467 | 0.6724 | 0.6799 | 0.6894 | 0.6990 | 0.7157 |
| BiRW | 0.3872 | 0.5957 | 0.6468 | 0.6822 | 0.7071 | 0.7246 |
| BiGCN | 0.2789<sup>★</sup> | 0.3523<sup>★</sup> | 0.3972<sup>★</sup> | 0.4493<sup>★</sup> | 0.5124<sup>★</sup> | 0.5847<sup>◇</sup> |
| GCN Cell | 0.2764<sup>★</sup> | 0.3537<sup>◇</sup> | 0.3997<sup>◇</sup> | 0.4554<sup>★</sup> | 0.5179<sup>★</sup> | 0.5935<sup>★</sup> |
| GCN Gene | 0.2833 | 0.3637<sup>★</sup> | 0.4005<sup>★</sup> | 0.4500<sup>◇</sup> | 0.5127<sup>◇</sup> | 0.5953 |

**Table 6.** Correlations on 5k cell simulation data (red, orange and blue indicate the top three methods).

| Dropout Ratio | 10% | 40% | 50% | 60% | 70% | 80% |
| --- | --- | --- | --- | --- | --- | --- |
| Density | 30.37 | 20.25 | 16.88 | 13.51 | 10.13 | 6.75 |
| ALRA | 0.4244 | 0.3741 | 0.3822 | 0.3612 | 0.3503 | 0.2392 |
| MAGIC | 0.7357<sup>★</sup> | 0.7201<sup>★</sup> | 0.7102<sup>★</sup> | 0.6956<sup>◇</sup> | 0.6743 | 0.6474<sup>★</sup> |
| DeepImpute | 0.6518 | 0.5244 | 0.5237 | 0.5142 | 0.5318 | 0.5403 |
| GNNImpute | 0.2664 | 0.2263 | 0.2185 | 0.2050 | 0.1943 | 0.1818 |
| BiRW | 0.6633 | 0.6101 | 0.5808 | 0.5447 | 0.4840 | 0.2905 |
| BiGCN | 0.7156<sup>★</sup> | 0.7085<sup>◇</sup> | 0.7039<sup>◇</sup> | 0.6991<sup>★</sup> | 0.6872<sup>★</sup> | 0.6694<sup>★</sup> |
| GCN Cell | 0.7175<sup>◇</sup> | 0.7072<sup>★</sup> | 0.6983 | 0.6890 | 0.6813<sup>★</sup> | 0.6430<sup>★</sup> |
| GCN Gene | 0.7103 | 0.7040 | 0.7007<sup>★</sup> | 0.6953<sup>★</sup> | 0.6843<sup>◇</sup> | 0.6558<sup>◇</sup> |

Finally, the results on the 10k cell simulation data, where the number of cells exceeds the number of genes, are shown in Table 7 and Table 8. Again, BiGCN has the lowest MSE for four datasets and the second lowest for two datasets. GCN Cell and GCN Gene also consistently show low values, indicating that GCN captures these features well. In the correlation results, BiGCN achieves the best results in three datasets, the second highest in two datasets, and the third highest for one.

**Table 7.** MSE on 10k cell simulation data (red, orange and blue indicate the top three methods).

| Dropout Ratio | 10% | 40% | 50% | 60% | 70% | 80% |
| --- | --- | --- | --- | --- | --- | --- |
| Density | 30.25 | 20.17 | 16.81 | 13.45 | 10.08 | 6.72 |
| ALRA | 2.2104 | 3.2241 | 3.7769 | 4.4294 | 5.5796 | 7.5676 |
| MAGIC | 0.2624<sup>*</sup> | 0.3620 | 0.4154 | 0.4775 | 0.5463 | 0.6165 |
| DeepImpute | 0.9604 | 0.8388 | 0.7669 | 0.7753 | 0.6317 | 0.5657<sup>*</sup> |
| GNNImpute | 0.6386 | 0.6652 | 0.6726 | 0.6808 | 0.6906 | 0.7016 |
| BiRW | 0.3835 | 0.5884 | 0.6384 | 0.6734 | 0.6984 | 0.7169 |
| BiGCN | 0.2732<sup>◇</sup> | 0.3492<sup>*</sup> | 0.3912<sup>*</sup> | 0.4452<sup>*</sup> | 0.5043<sup>*</sup> | 0.5777<sup>◇</sup> |
| GCN Cell | 0.2734<sup>◇</sup> | 0.3498<sup>◇</sup> | 0.3915<sup>◇</sup> | 0.4452<sup>◇</sup> | 0.5076<sup>◇</sup> | 0.5824<sup>◇</sup> |
| GCN Gene | 0.2828 | 0.3611<sup>◇</sup> | 0.4008<sup>◇</sup> | 0.4523<sup>◇</sup> | 0.5137<sup>◇</sup> | 0.5980 |

**Table 8.** Correlations on 10k cell simulation data (red, orange and blue indicate the top three methods).

| Dropout Ratio | 10% | 40% | 50% | 60% | 70% | 80% |
| --- | --- | --- | --- | --- | --- | --- |
| Density | 30.25 | 20.17 | 16.81 | 13.45 | 10.08 | 6.72 |
| ALRA | 0.4227 | 0.3889 | 0.3844 | 0.2659 | 0.2839 | 0.3322 |
| MAGIC | 0.7388<sup>*</sup> | 0.7223<sup>*</sup> | 0.7129<sup>*</sup> | 0.6998<sup>◇</sup> | 0.6806<sup>◇</sup> | 0.6538<sup>◇</sup> |
| DeepImpute | 0.6604 | 0.5269 | 0.5190 | 0.5177 | 0.5339 | 0.5420 |
| GNNImpute | 0.2762 | 0.2323 | 0.2250 | 0.2163 | 0.2032 | 0.1896 |
| BiRW | 0.6654 | 0.6128 | 0.5840 | 0.5519 | 0.5121 | 0.3549 |
| BiGCN | 0.7196<sup>◇</sup> | 0.7110<sup>◇</sup> | 0.7064<sup>◇</sup> | 0.7030<sup>*</sup> | 0.6986<sup>*</sup> | 0.6848<sup>*</sup> |
| GCN Cell | 0.7190<sup>◇</sup> | 0.7104<sup>◇</sup> | 0.7091<sup>◇</sup> | 0.7019<sup>◇</sup> | 0.6984<sup>◇</sup> | 0.6747<sup>◇</sup> |
| GCN Gene | 0.7078 | 0.6981 | 0.6926 | 0.6922 | 0.6784 | 0.6349 |

In summary, MAGIC generally performs better in high density data (e.g. 10% dropout) since MAGIC uses the k-nearest neighbor method (KNN), which works well on dense data. Since BiGCN uses linear kernels, which capture global information about all cells, its imputations are significantly better on sparse data. In addition, DeepImpute may show low MSE for extremely sparse data. This occurs because the loss function of DeepImpute exclusively learns the nonzero entries and produces extremely low values for sparse data with few nonzero entries, leading to overfitting and resulting in low correlations. BiGCN emerges as the most accurate and stable method when the dropout ranges from 40-80%, a significant finding given that real scRNA-seq data is often impacted by dropout rates of 10-80% Stegle et al. [2015]. The complete results of more dropout rates are shown in the supplementary document.

### Imputation results of PBMC scRNAseq data with aggregated bulk reference

Since ground truth data is only available for evaluation in simulation studies, the evaluation on the real scRNA-seq data is much more difficult. This is mainly due to the ambiguity between true biological zeros and the technical zeros (dropouts) in the real data. Standard cross-validation on all the values or only the non-zero entries is an insufficient evaluation. Additionally, regarding clustering, many cell classes associated with scRNA-seq are annotated using tools like Seurat or spectral clustering. Therefore, the appropriateness of evaluating clustering performance on these datasets using metrics such as ARI is also ambiguous.

To avoid some of these pitfalls, we created an aggregated bulk reference dataset derived from the PBMC dataset. The imputation methods do not observe the data used to create the aggregated bulk reference; instead, they are solely provided with the 10,000 cells for which the bulk reference was created in relation to. Comparing the imputation results on these 10,000 cells to the aggregated bulk reference allows us to assess an imputation method’s capability to recover true biological zeros and dropout. Table 9 assesses the imputation performance of each method using the MSE for nonzero entries, the MSE for zero entries, and the correlations for nonzero entries. BiGCN achieved the highest scores across all data and metrics, except for the MSE of nonzero entries with 30 neighbors, which might be due to the over-smoothing with the large number of neighbors. These results strongly support that BiGCN accurately imputes real scRNA-seq data and suggest that it can identify true biological signals and distinguish true biological zeros from dropouts.

**Table 9.**
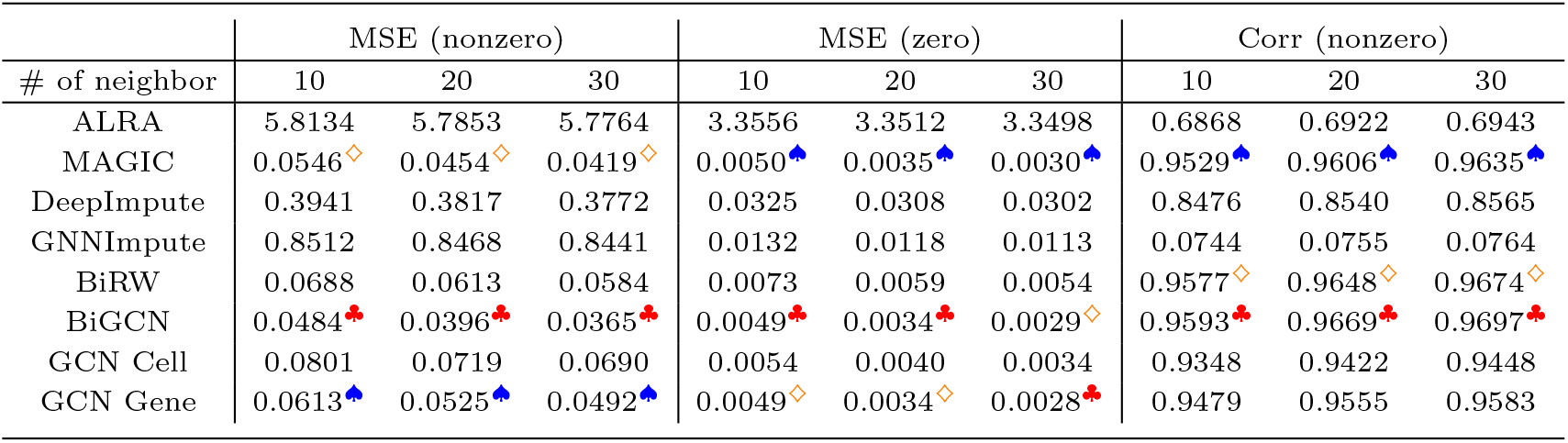
MSE of zero entries, MSE of zero entries and correlation of nonzero entries in PBMC data.

| # of neighbor | MSE (nonzero) |  |  | MSE (zero) |  |  | Corr (nonzero) |  |  |
| --- | --- | --- | --- | --- | --- | --- | --- | --- | --- |
|  | 10 | 20 | 30 | 10 | 20 | 30 | 10 | 20 | 30 |
| ALRA | 5.8134 | 5.7853 | 5.7764 | 3.3556 | 3.3512 | 3.3498 | 0.6868 | 0.6922 | 0.6943 |
| MAGIC | 0.0546<sup>◇</sup> | 0.0454<sup>◇</sup> | 0.0419<sup>◇</sup> | 0.0050<sup>◇</sup> | 0.0035<sup>◇</sup> | 0.0030<sup>◇</sup> | 0.9529<sup>◇</sup> | 0.9606<sup>◇</sup> | 0.9635<sup>◇</sup> |
| DeepImpute | 0.3941 | 0.3817 | 0.3772 | 0.0325 | 0.0308 | 0.0302 | 0.8476 | 0.8540 | 0.8565 |
| GNNImpute | 0.8512 | 0.8468 | 0.8441 | 0.0132 | 0.0118 | 0.0113 | 0.0744 | 0.0755 | 0.0764 |
| BiRW | 0.0688 | 0.0613 | 0.0584 | 0.0073 | 0.0059 | 0.0054 | 0.9577<sup>◇</sup> | 0.9648<sup>◇</sup> | 0.9674<sup>◇</sup> |
| BiGCN | 0.0484<sup>*</sup> | 0.0396<sup>*</sup> | 0.0365<sup>*</sup> | 0.0049<sup>*</sup> | 0.0034<sup>*</sup> | 0.0029<sup>◇</sup> | 0.9593<sup>*</sup> | 0.9669<sup>*</sup> | 0.9697<sup>*</sup> |
| GCN Cell | 0.0801 | 0.0719 | 0.0690 | 0.0054 | 0.0040 | 0.0034 | 0.9348 | 0.9422 | 0.9448 |
| GCN Gene | 0.0613<sup>◇</sup> | 0.0525<sup>◇</sup> | 0.0492<sup>◇</sup> | 0.0049<sup>◇</sup> | 0.0034<sup>◇</sup> | 0.0028<sup>*</sup> | 0.9479 | 0.9555 | 0.9583 |

Discovering novel methods for assessing imputation is crucial, and further improvements still need to be made. While this experiment yielded successful imputation and cell clustering, it also raised numerous new questions. We would like to address these questions in future experiments.

### Clustering results on 10 scRNAseq datasets

Next, clustering results are shown in Fig 2. Note that for the clustering tasks BiGCN, GCN Cell, and GCN Gene use normalized imputation (see Methods). BiGCN achieves the highest ARI on six datasets and ARIs nearly equivalent to the highest in the remaining four datasets. Among the four datasets, ALRA has the highest ARI on two datasets, while MAGIC and Raw data exhibit the highest ARI in one dataset. GCN Cell, relying solely on cell information, generally also produces high results in cell clustering. The fact that BiGCN achieves the highest ARI on the majority of datasets suggests that considering gene co-expression during imputation enhances the signal used to distinguish cell types. However, the results of GCN Gene underperform GCN Cell, indicating that approaches like GCN Cell, which directly capture cell-cell similarity, yield imputations more advantageous for clustering.

**Fig. 2.**
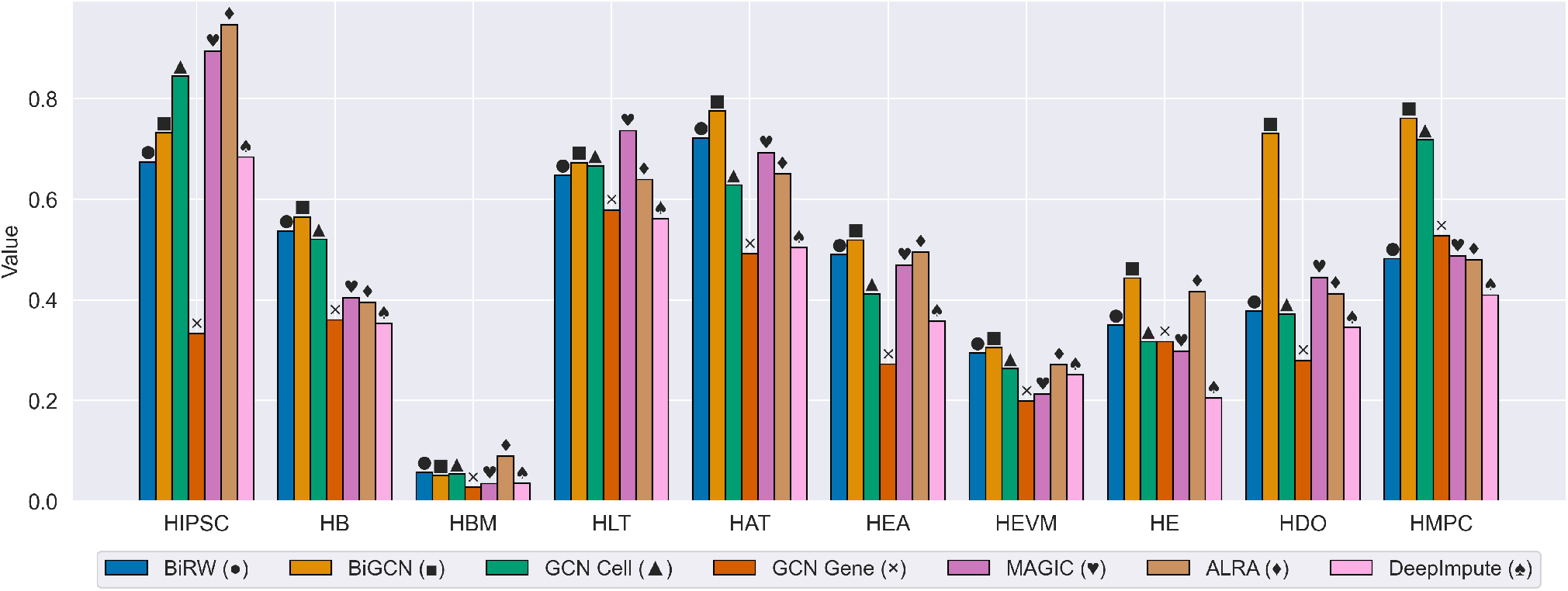
ARI for ten scRNAseq datasets.

Across five of the datasets, GCN Cell consistently outperforms MAGIC. While MAGIC’s imputation is based on aggregation of neighborhood data through KNN graphs, GCN Cell determines the imputation through graph convolution. Similar our simulation results, MAGIC tends to perform better on dense data, whereas GCN Cell is more effective on sparse data.

While ALRA achieves the highest ARI of the two datasets, ALRA performed very poorly in the imputation experiments, indicating ALRA might not be suitable for direct imputation. BiGCN significantly outperforms BiRW, which suggests the need for more advanced graph convolution as opposed to simple random walks to aggregate graph neighborhood information. We also attempted to visualize the results using UMAP and t-SNE, as shown in Figures S1 and S2. This visualization supports our claim that our method achieves better cluster separation.

### Runtime comparison

Figure 3 shows the execution times of ALRA, DeepImpute, MAGIC, BiRW, BiGCN, GCN Cell, GCN Gene and GNNImpute. The evaluation is conduced on an NVIDIA GeForce RTX 2080 Ti GPU with 12GB of VRAM.

**Fig. 3.**
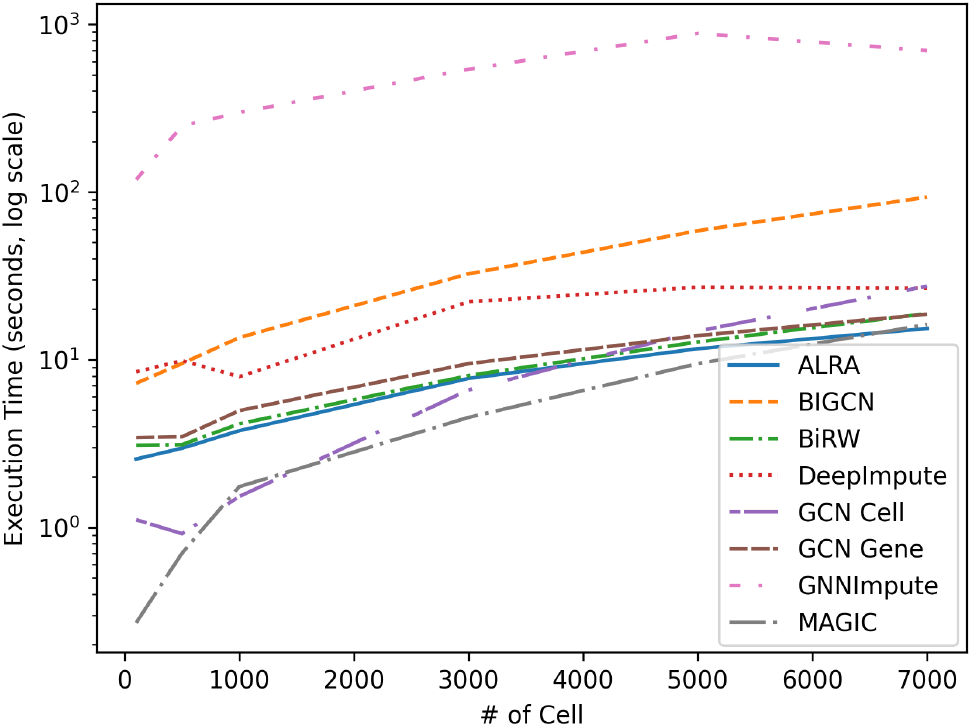
Runtime comparison. The y-axis indicates the execution time on a logarithmic scale, and the horizontal axis indicates the number of cells.

First, MAGIC, ALRA, and BiRW are based on a calculation of similarities and learning statistical models, and they are fast even when the number of cells is large. In deep learning-based methods, both GCN Cell and GCN Gene operate quickly. The cell size clearly influences GCN Cell speed with longer runtime as the number of cells increases. Conversely, the speed of GCN Gene is determined by the number of genes, and its performance remains relatively stable even as the number of cells increases. DeepImpute is a standard feed-forward neural network that runs very efficiently. BiGCN is slightly slower than its counterparts, as it depdents on both the number of cells and genes. Notably, BiGCN still completes within 100 seconds when using 7000 cells. In addition, GNNImpute is a more complex model utilizing a Graph Attention Network, which significantly increases the number of parameters compared to BiGCN and substantially increases the runtime. In addition, the execution time of scGNN2.0 is reported to be overwhelmingly slower with approximately 100 seconds per epoch for just 400 cells [Gu et al., 2022]. Based on the above results, we conclude that BiGCN can scale to large problem sizes on real datasets, where other more complex models may encounter difficulties.

## Conclusions

In this work, we developed a new method, BiGCN, which effectively utilizes both cell similarities and gene co-expression information to impute scRNA-seq data. In our experiments, we evaluated the impact of combining these two types of information and demonstrated that BiGCN represents a significant improvement over existing imputation methods. The comparison of GCN Cell, GCN Gene, and BiGCN shows the importance of integrating both gene-gene and cell-cell information. Surprisingly, both GCN Cell and GCN Gene produce imputations that are informative for clustering, indicating that capturing characteristics of gene co-expression is nearly as beneficial as capturing cell similarities.. BiGCN, which combines these two sources of information, works the best. The comparisons between GCN Cell and MAGIC, as well as between BiGCN and BiRW, also demonstrate that more complex graph convolution models like GCN can more effectively capture cell and gene similarities compared to a simple random walk.

One interesting observation in this study is the “paradoxical” relationship between imputation accuracy by expression value and capturing results. For instance, MAGIC performs well at imputation but generates unsatisfactory clustering results. On the other hand, ALRA shows poor imputation results but accurate clustering performance. This is also observed in the application of BiGCN. To mitigate this issue, we introduced the variation “imputation for normalized expression.” just for clustering. This variation allows for imputation to preserve more cell characteristics.

## APPENDIX A

**Fig. S1.**
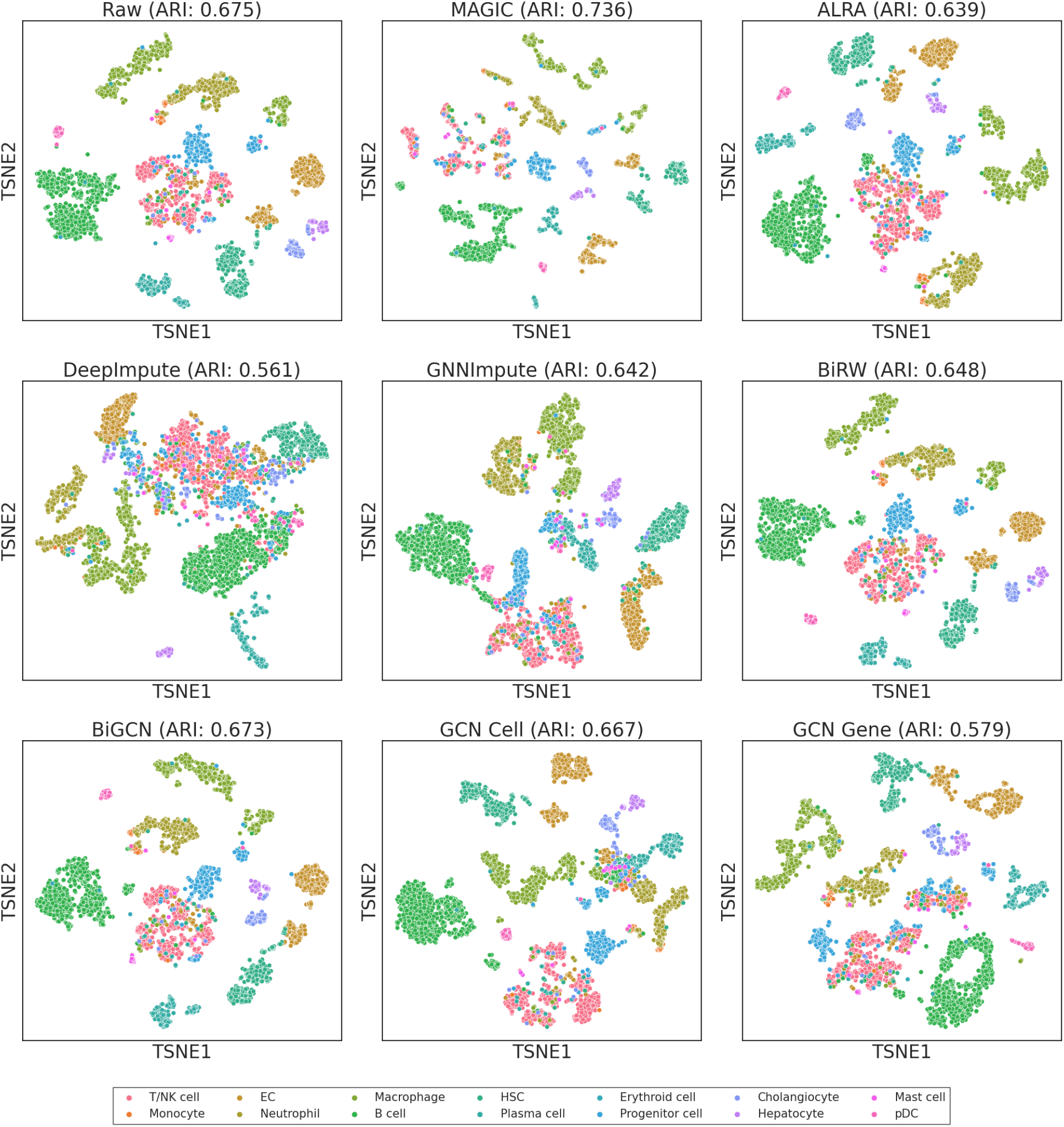
Visualization of HLT dataset with UMAP.

**Fig. S2.**
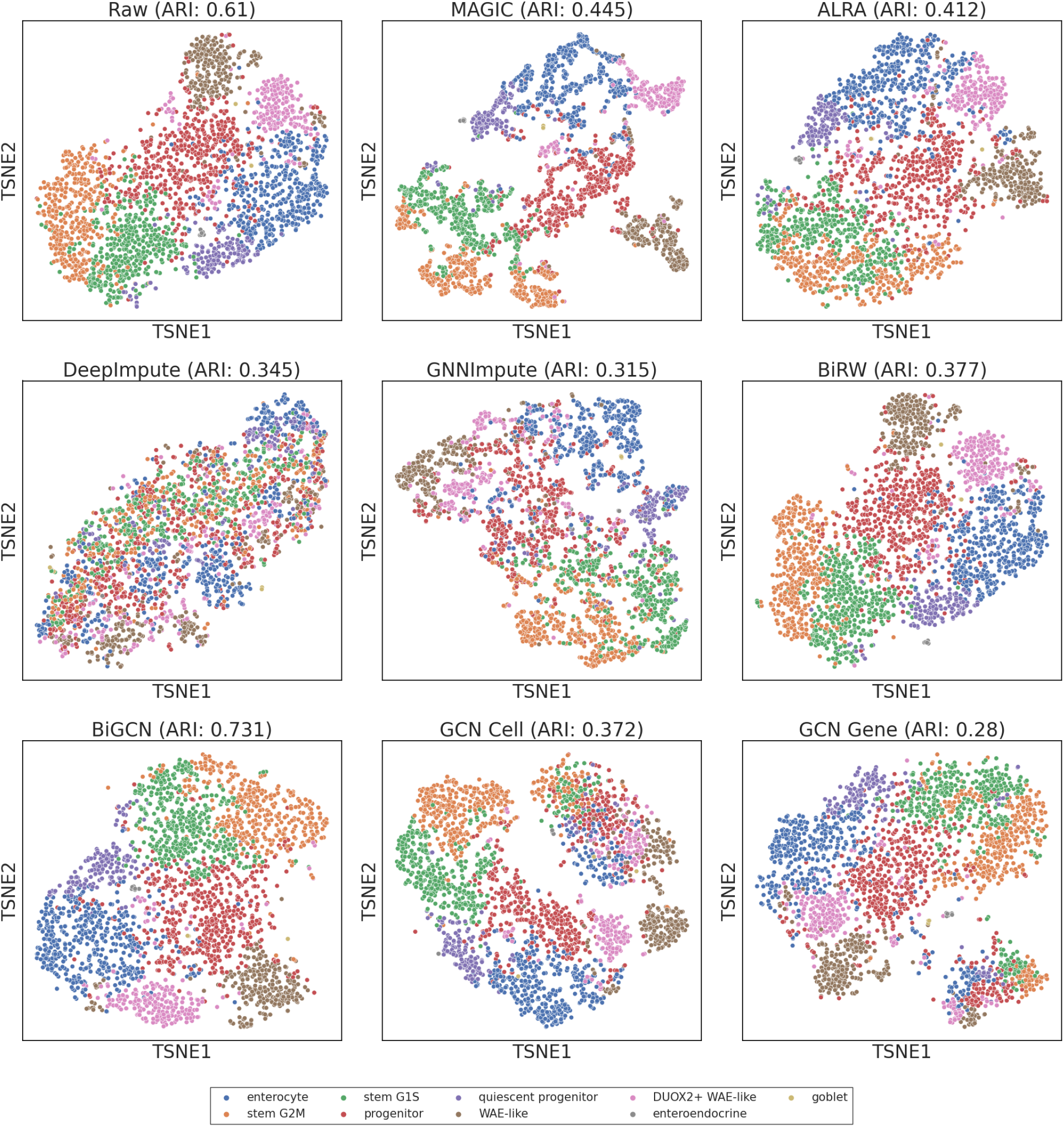
A visualization of the data imputation for HDO data using UMAP.

